# The influence of power-training and aging on endomysium content and fiber cross-sectional area in the human soleus muscle

**DOI:** 10.1101/2025.08.28.672902

**Authors:** Christoph S. Clemen, Sebastian W. Humbsch, Carolin Berwanger, Leonid Mill, Andreas Schmidt, Rolf Schröder, Jörn Rittweger

## Abstract

The Master Athletic Laboratory Study of Intramuscular Connective Tissue (MALICoT, DRKS00015764) set out to analyze the endomysium content of the human soleus muscle in response to athletic exercise and aging. Forty-three healthy male study participants were grouped into young (20–35 years) non-physically active controls (n=12), young power-trained athletes (n=10), older (60–75 years) non-physically active controls (n=11), and older power-trained athletes (n=10). A single biopsy was taken from the left soleus muscle of each participant, and cryo-sections were used for i) routine histological staining and myopathological evaluation, ii) deep learning-based image analysis of the H&E- and MHC-stained sections, iii) laminin-*γ*-1/collagen IV double- and collagen I and III single-immunofluorescence staining, and iv) quantitative proteomic analysis. Examiner-based myopathological evaluation revealed normal skeletal muscle in 26 participants, while 11, 4, 1 and 1 biopsies showed unspecific myopathological changes, chronic neurogenic atrophy, type II fiber atrophy, and unspecific myositic changes, respectively. Analysis of the H&E- and MHC-stained sections as well as the laminin-*γ*-1/collagen I-, collagen III- and collagen IV-immunostained sections revealed an approximately 1.3-fold increase in the ‘mean fiber area’ in response to power-training in young participants and aging in unathletic participants. No significant change was detected in endomysium thickness or area. Furthermore, proteomic analysis did not reveal any group-specific change except for plasma membrane calcium-transporting ATPase 2 being less abundant in the soleus muscles of aged power-trained athletes. Overall, the data show that neither athletic exercise nor age significantly affected the content or composition of the endomysium in human soleus muscle tissue.

**New & Noteworthy:** MALICoT (DRKS00015764) set out to analyze the endomysium of soleus muscle in response to athletic exercise and aging. Myopathological evaluation of biopsies from 43 asymptomatic male study participants revealed normal skeletal muscle in only 26 of them. Neither exercise nor age significantly affected the endomysium content or proteomic profiles of human soleus muscle tissue. However, power-training in young participants and aging in unathletic participants were associated with significant increases in soleus muscle fiber cross-sectional area.

## Introduction

Skeletal muscle function declines with age, leading to a reduction in muscle mass and mechanical output. Although numerous studies have investigated this decline in muscle mass and function (Larsson et al., 2019), the intramuscular connective tissue has received little attention. The endomysium and perimysium are generally thought to play a mechanical role, both in distributing mechanical stresses and allowing slippage between fibers and between fascicles (Purslow, 2020). It has been assumed that aging is associated with the accumulation of intramuscular connective tissue in mice, rats and humans (Alnaqeeb et al., 1984; Fede et al., 2022; Yeung et al., 2023). We previously reported that the endomysium content in the soleus muscles of participants increased in response to 60 days of experimental bed rest when normalized to muscle fiber area, but not when normalized to muscle fiber number (Thot et al., 2021), suggesting that the total intramuscular connective tissue content remains constant during at least two months of immobilization. Thus, it is possible that the reported accumulation of intramuscular connective tissue with age is more an effect of fiber shrinkage rather than a true increase in intramuscular connective tissue content.

In the Master Athletic Laboratory Study of Intramuscular Connective Tissue (MALICoT), which was conducted as a complementary study to the research on bed rest-induced muscle wasting (Thot et al., 2021), we addressed the hypothesis that aging and training status would both affect the endomysium content in the soleus muscle. Since aging is typically associated with a more sedentary lifestyle (Varo et al., 2003), so that the muscle phenotype observed in older individuals is influenced by both senescence and immobilization, we focused on Masters athletes. These athletes continue to train and compete in sport at older ages, often well into their 8^th^ or 9^th^ decade of life (Tanaka et al., 2019). We further focused on the soleus muscle, which acts as a spring muscle during running (Lai et al., 2019), and selected athletes specialized in sprinting and jumping, as elastic energy storage in the calf muscles is paramount in these activities (Cavagna et al., 2008). We assumed that the mechanical demands on the intramuscular connective tissue would be greatest during spring action. Due to the intrinsic difficulty of organizing a longitudinal, interventional study in healthy human volunteers, we chose a fully factorial cross-sectional approach comparing four groups of athletes and non-athletes at young and old ages.

## Materials and Methods

### Ethical approval and study design

The Master Athletic Laboratory Study of Intramuscular Connective Tissue (MALICoT) was approved by the Ethics Committee of the North Rhine Medical Association, Düsseldorf, Germany, reference number 2018269, and was registered in the German Clinical Trials Register, DRKS-ID DRKS00015764. All participants gave written informed consent before participating in the study. The study adhered to the principles of the Declaration of Helsinki and to the General Data Protection Regulation (GDPR).

The start of recruitment was hampered by the beginning of the COVID-19 pandemic. This led to testing sessions being skipped or postponed due to containment measures or COVID infection of staff or participants. As a result, the recruitment aim of 12 participants per group (48 in total) could not be reached. In the end, 43 healthy male participants with a body mass index ≤28 kg/m^2^ were included in the study and tested between 5^th^ August 2020 and 26^th^ May 2021. Blinded to the investigators, the participants were divided into four different groups, young non-athletes (n=12) and young power-trained athletes (n=10) (age range 20 to 35 years) and old non-athletes (n=11) and old power-trained athletes (n=10) (age range 60 to 75 years). The participants completed a questionnaire to quantify their exercise-related metabolic equivalents for task (METs) (Frey et al., 1999), underwent medical examination, and anthropometric data were collected. Participants who did at least four hours of running training per week, which was also monitored by means of an accelerometric recording box worn for at least one week, and participated in running competitions were categorized as athletes. Participants were categorized as unathletic if they had little or no sports-related physical activity, defined by an activity-related energy expenditure of ≤2 METs. Further information on MALICoT has been published, including participants characteristics and biomechanical parameters (Sanchez-Trigo et al., 2022), bone geometry data (Scorcelletti et al., 2023), physical activity and cardiometabolic risk profiling data (Rittweger et al., 2024), and muscle volume, fat fraction and power data (Zange et al., 2025).

### Soleus muscle biopsy

Biopsies were taken from the left soleus muscle of all participants under local anesthesia and in sterile conditions using a vacuum-assisted biopsy system with a 10G needle (Vacora Biopsy System type VF2019, Bard). This yielded cylindrical muscle tissue specimens that were largely mechanically intact, measuring approximately 19 mm in length, 3 mm in diameter and 150 mg in weight. The physical principle of this automated biopsy system with single-use biopsy needles is similar to the manual Bergström biopsy technique, but with inverted tubes (the outer one is mobile) and with much less mechanical damage to the biopsy specimen (Akarolo-Anthony et al., 2012; Lee et al., 2020). Part of the tissue cylinders were mounted on cork plates (15 mm diameter, 3 mm thickness) with fiber orientation for transverse cryosectioning, covered with Tissue-Tek OCT Compound (Sakura Finetek, Torrance, CA, USA), snap-frozen in liquid nitrogen-cooled isopentane to avoid freezing artefacts, collected in 5 ml LDPE sample vials with caps, and stored at −80°C for subsequent analyses.

### Histological staining and myopathological evaluation

H&E, Oil red O, Gömöri trichrome, COX/SDH, PAS, slow MHC (MHCs), fast MHC (MHCf), developmental MHC (MHCd) and neonatal MHC (MHCn)-stained cross-sections of 6 µm thickness of the soleus muscle biopsy specimens were prepared with a cryotome (Leica CM 1850 UV) and used for histopathological evaluation by an experienced myopathologist.

### Parameter acquisition using an automated deep learning-based biomedical image approach

Whole slide images of the H&E-, MHCs- and MHCf-stained soleus muscle sections (typically three sections per slide) were recorded by a slide scanner (NanoZoomer S60, Hamamatsu, Japan) and subjected to an automated deep learning-based image analysis software tool based on photorealistic, computer graphics generated synthetic data in muscle histopathology (Mill et al., 2025). In this study, the software was used to identify the stained muscle tissue sections, transverse muscle fiber orientation, sectioning and staining artefacts, and to segment muscle fibers (H&E, MHCs and MHCf stains) and endomysium (H&E stain only). For H&E-stains, the following parameters were obtained for each muscle fiber, each manually added region of interest (ROI), which typically contained no perimysium and about 100 muscle fibers (sufficient for reliable quantification (McCall et al., 1998)), each whole muscle tissue section, and each full slide, and output as individual, total or mean values: ‘number of samples’, ‘tissue area’, ‘number of fibers’, ‘fiber area’, ‘fiber area/tissue area’, ‘fiber diameter’, ‘fiber perimeter’, ‘endomysium area’, ‘endomysium area/tissue area’, ‘endomysium thickness’, ‘endomysium area/fiber area’, and ‘endomysium area/fiber number’. Note that setting a ROI in this software tool does not result in fibers being trimmed at the edges of a ROI as seen in figure 3. In the case of MHCs- and MHCf-stains, for each muscle fiber, each whole muscle tissue section and each full slide, the individual, total or mean values ‘number of samples’, ‘number of fibers’, ‘MHC-staining positivity’, ‘fiber area’, and ‘fiber diameter’ were determined.

### Indirect immunofluorescence staining

Cryostat sections of 6 µm thickness were collected on microscope slides, air-dried for 30 min, fixed in formaldehyde prepared from 0.5% paraformaldehyde for 10 min, washed three times in PBS for 5 min, and blocked with 5% NGS/0.2% Tween-20 in PBS for 30 min (laminin-*γ*-1/collagen IV double-staining) or 10% NGS/0.2% Tween-20/0.15% glycine for 3h (collagen I and collagen III single staining). For staining of plasma membrane calcium-transporting ATPase 2 (ATP2B2/PMCA2), dried sections were fixed in −20°C cold acetone for 10 min, washed and blocked with 5% NGS/0.2% Tween-20 in PBS. Blocked sections were incubated with the primary antibodies diluted in the respective blocking buffer at room temperature for 1h, for laminin-*γ*-1/collagen IV double-staining with rabbit polyclonal anti-laminin-*γ*-1 (Immundiagnostik, AP1001.2, 1:200) and mouse monoclonal anti-collagen IV (Abcam, AB86042, 1:100), for the collagen I and collagen III single stains with either mouse monoclonal anti-collagen I (Sigma-Aldrich, C2456, 1:100) or mouse monoclonal anti-collagen III (Sigma-Aldrich/Merck-Millipore, MAB3392, 1:100), and for plasma membrane calcium-transporting ATPase 2 with rabbit polyclonal anti-PMCA2 ATPase (Invitrogen, PA1-915, 1:200). After six washes in PBS for 10 min each, sections were incubated with the appropriate secondary antibodies diluted 1:500 in blocking solution, goat anti-rabbit AlexaFluor568 (Invitrogen, A-11011) and goat anti-mouse AlexaFluor647 (Invitrogen, A-21236) together with DAPI at 1:1000 dilution for 45 min; for PMCA2 goat anti-rabbit AlexaFluor555 (Invitrogen, A-21429) diluted 1:400. Final washes were three times with 0.5% Triton X-100 in PBS and five times with PBS for 15 min each, before sections were rinsed in ddH_2_O and embedded in Mowiol/DABCO.

### Parameter acquisition by microscope and image software and manual image analysis

Laminin and collagen immunofluorescence images were recorded using a Zeiss Axio Imager.Z2m microscope (Carl Zeiss Microscopy, Oberkochen, Germany) with a 40x oil objective (NA 1.4). The Zeiss Zen software version 3.4 was used to stitch together the tile images into full section overviews and to set a region of interest (ROI) for each tissue specimen for subsequent image analysis. A ROI typically contained 100 muscle fibers (sufficient for reliable quantification (McCall et al., 1998)) and was selected so that the muscle fibers had a clear transverse orientation in the section, without perimysium and without artefacts from cryosectioning or staining.

Based on a previously established protocol, in which the laminin-*γ*-1 stained basement membrane was used to segment muscle fibers using the Zeiss Zen software version 3.4 (Thot et al., 2021), here the fluorescence channels of laminin-*γ*-1 and collagen IV were merged, as this resulted in an improved segmentation pattern. The software-derived parameters of muscle fibers were number of fibers, fiber area, minimal and maximal Feret-diameters, fiber perimeter, and ROI dimensions. In addition, the number of trimmed fibers, i.e., the fibers at the edges of a ROI as seen in figure 2, was determined manually to correct the number of fibers by calculating the difference between the software-derived total number of fibers minus half of the number of trimmed fibers. The mean fiber area was calculated as the software-derived total fiber area, including the trimmed fibers, divided by the corrected number of fibers. As the endomysial space is delineated by the laminin-*γ*-1 and collagen IV stained basement membranes of adjacent muscle fibers, the endomysial parameters were calculated according to (Thot et al., 2021). The total endomysium area was the difference between the ROI area and the total fiber area, and the mean endomysium thickness adjusted for trimmed fiber effects was the total endomysium area divided by the difference of the half of the total fiber perimeter minus the ROI perimeter.

In the case of the analysis of the collagen I and collagen III stained muscle sections, in which the endomysial area is labelled, the respective ROI images and parameters were exported from the Zeiss Zen software, and the ROI images were imported for a pixel-based analysis into the freeware IrfanView version 4.60 (https://www.irfanview.com/) to derive endomysial and muscle fiber parameters. Imported ROI images were converted to black and white images, and the numbers of all ROI and black endomysial and white muscle fiber area pixels were obtained as well as pixel dimensions to calculate areas. The total number of fibers and trimmed fibers were counted manually and used to calculate the corrected number of fibers and the mean fiber area. The mean endomysial diameter was calculated as above, considering muscle fibers as squares, so that the total fiber perimeter was estimated as being four times the square root of the mean fiber area multiplied by the corrected number of fibers.

### Muscle parameters, statistical analysis, and miscellaneous methods

Four main muscle parameters were obtained or calculated for the purpose of this work using different image analysis approaches as described above, in order to determine the soleus muscle endomysium content as a main objective of MALICoT, and to simplify and improve approaches for quantitating muscle connective tissue. All data were collected in a structured manner in Excel tables (Excel 2016, Microsoft) and imported into GraphPad Prism version 10.5.0.774 (GraphPad Software, Boston, MA) to perform basic statistics, normality tests, unpaired two-sample Welch’s t-tests or Mann-Whitney U tests, as appropriate, and scatter plots for data visualization. The number of samples and p-values are given in the figure legends. Graphs and microscopic and other images were further processed and figures were assembled using CorelDraw Graphics Suite X7 (Corel Corporation, Ottawa, Canada).

### Proteomics

Human soleus muscle cryosections (10 sections á 20 µm thickness per sample) were dissolved in 10% SDS in PBS by boiling at 95°C for 10 min and subsequent sonication (30 s on, 30 s off) using a Bioruptor sonication bath at 14°C for 10 min. Disulfide bonds were reduced by addition of 5 mM DTT and incubation at 55°C for 30 min, and alkylated by addition of 40 mM chloroacetamide and incubation at room temperature for another 30 min. Samples were centrifuged at 20,000 x g for 10 min and 100 µL of the supernatant of each sample were transferred into a 96-well plate PCR plate. Magnetic SP3 protein affinity beads (2.5 µL) and immediately 100 µL of acetonitrile were added to the samples. Proteins were allowed to bind for 10 min, followed by washing with 70% ethanol (3x 100 µL) and 100% acetonitrile (100 µL). After drying the beads, proteins were digested with 0.5 µg trypsin in 50 mM ammonium bicarbonate at 35°C for 12h. Subsequently, peptide mixtures were desalted on SDB-RP stage tips, vacuum dried and redissolved in LC-MS loading buffer (5% FA, 2% acetonitrile in water).

For LC-MS analysis, 300 ng of peptides were separated using a nanoElute HPLC chromatography system with a 5 mm trapping column and a 25 cm separation column. A linear gradient from 5 to 35% acetonitrile was applied over a time of 35 min to fractionate the peptide sample (pulled tip, 300 nL/min, 50 min). Eluting peptides were directly ionized and detected in a timsTOF Pro2 mass spectrometer using a data-independent acquisition strategy (12×50 m/z Window 4 IMS PASEF-DIA method). Data files were searched against the Swissprot/Uniprot human protein database using the DIA-NN 1.8.1 (Data-Independent Acquisition by Neural Networks) software suite. Search settings were fixed modification for Cys carbamidomethylation, variable oxidation of methionine side chains, sample dependent mass accuracy, double pass mode for protein identification, precursor m/z 400-1100, fragment m/z 250-1700, report of protein identification quality and heuristic protein inference. Protein abundances were recovered from proteotypic peptides using an in-house R-script based on the one supplied by DIA-NN. Sample statistics were performed in Perseus. Raw data have been deposited to the ProteomeXchange Consortium (Deutsch et al., 2023) via the PRIDE (Perez-Riverol et al., 2022) partner repository with the dataset identifier PXD070244.

### PMCA2 immunofluorescence microscopy and image processing

Plasma membrane calcium-transporting ATPase 2 (PMCA2) immunofluorescence images were recorded using an Infinity Line system (Abberior Instruments GmbH, Göttingen, Germany) with a UPLXAPO60XO NA 1.42 objective and Imspector software version 16.3.16100 in LightBox mode. The confocal images were further deconvolved using Huygens Essential version 23.04.0p6 (Scientific Volume Imaging B.V., Hilversum, The Netherlands).

## Results

### The spectrum of myopathological findings in young and aged healthy males

Cryo-sections of the soleus muscle biopsy specimens from all 43 clinically healthy study participants were used for a myopathological evaluation based on hematoxylin and eosin (H&E), Oil red O, Gömöri trichrome (GT), combined cytochrome c oxidase/succinate dehydrogenase (COX/SDH), periodic acid Schiff (PAS), and myosin heavy chain (MHC) isoform staining (Fig. 1). This examiner-based myopathological evaluation denoted the diagnosis of ‘normal skeletal muscle tissue’ in 26 participants, in 10 out of 12 biopsies from young unathletic participants (red group in Figs. 1,6), in 9 out of 10 young power-trained athletes (blue group in Figs. 1,6), in 7 out of 11 aged unathletic participants (purple group in Figs. 1,6), but in none of the 10 aged power-trained athletes (green group in Figs. 1,6). Three biopsies of the latter group and one more of the group of aged unathletic participants showed signs of chronic neurogenic atrophy (Fig. 1; and red symbols in the scatter plots of Fig. 4) defined by the presence of type I and type II fiber grouping along with atrophic fibers and nuclear clumps. The group of aged power-trained athletes had two more biopsies with pathological findings, one case of type II fiber atrophy (Fig. 1; and magenta symbols in the scatter plots of Fig. 4) defined by a large number of type II fibers with a >12 % smaller fiber diameter compared to type I fibers, and one case of unspecific myositic changes (Fig. 1; and turquoise symbols in the scatter plots of Fig. 4) defined by a small endomysial and small perimysial inflammatory infiltrate, groups of atrophic fibers, sporadic nuclear clumps, and an increased number of fibers with centralised nuclei. In additional 11 biopsies (3 from young and 8 from aged participants) (Fig. 1; and green symbols in the scatter plots of Fig. 4) unspecific myopathological changes defined by minor pathological alterations, i.e., sporadic atrophic fibers, were observed, which precluded the diagnosis of normal skeletal muscle.

**Figure 1.**
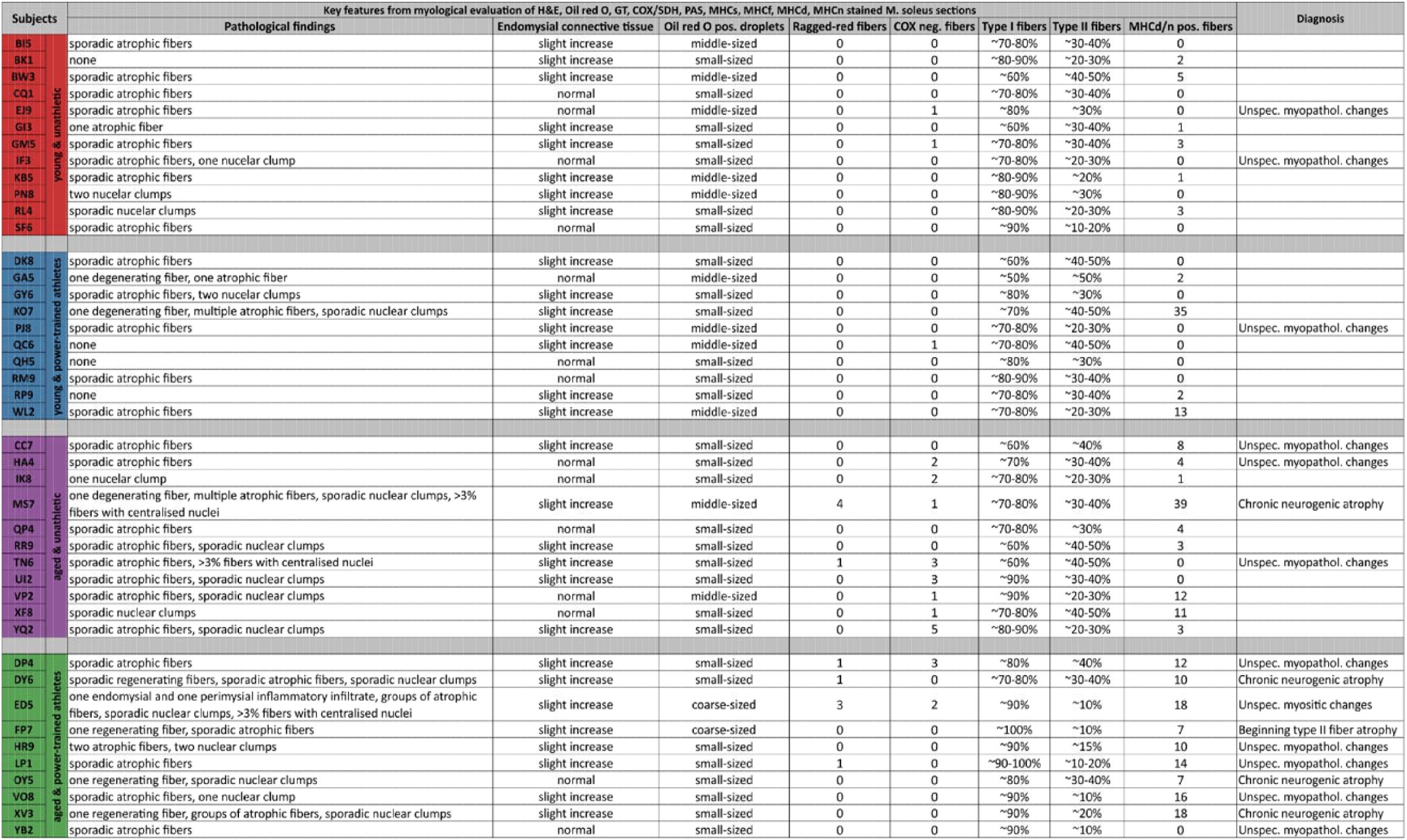
The spectrum of myopathological findings in the MALICoT. The Master Athletic Laboratory Study of Intramuscular Connective Tissue (MALICoT, registration number DRKS00015764) included a total of 43 participants, divided into 12 young (20 to 35 years) unathletic participants, 10 young power-trained athletes, 11 aged (60 to 75 years) unathletic participants and 10 aged power-trained athletes. Myopathological evaluation was based on H&E, Oil red O, Gömöri trichrome (GT), COX/SDH, PAS, slow MHC, fast MHC, developmental MHC and neonatal MHC stained cryo-sections of soleus muscle biopsy specimens. Pathological findings, specific myopathological findings; endomysial connective tissue, endomysial thickness estimated in the H&E-stained section, normal or slightly increased; Oil red O-positive droplets, classification as small-, middle-, or coarse-sized; ragged-red fibers, number of fibers containing clumps of mitochondria in the GT-stained section; COX-negative fibers, number of COX-negative fibers counted in the COX/SDH double-stained fibers; type I and type II fibers, estimated fractions in percent of fibers positive for either slow or fast MHC isoforms, fractions were determined in separately stained sections and therefore do not necessarily add up to 100%; MHCd/n positive fibers, total number of positive fibers from sections stained separately for developmental or neonatal MHC isoform expression; diagnosis, myopathological diagnosis, if present.

### Power-training in young participants and aging in unathletic participants: increase in the mean fiber cross-sectional area, but no significant change in endomysium content

To address the possible presence of more subtle changes in the amount or thickness of the endomysial connective tissue, different quantitative approaches were used to measure the endomysial content. Double immunofluorescence staining for laminin-*γ*-1 and collagen IV in transversal cryo-sections as described previously (Thot et al., 2021) was used to indirectly determine the endomysial content based on the lines of laminin-*γ*-1- and collagen IV-positive basal laminae and the non-stained stripe in between (Fig. 2A,B,G,H). In addition, collagen I (Fig. 2C,D,I,J) and collagen III (Fig. 2E,F,K,L) single immunofluorescence staining was used to directly visualize the connective tissue between the muscle fibers. As a new approach, an automated deep learning and photorealistic computer graphics generated synthetic data based artificial intelligence analysis tool (Mill et al., 2025) was used on images of H&E-stained sections (Fig. 3E-G,L-N). All analysis approaches were performed in a region of interest (ROI)-based manner, with the latter approach additionally performed as a full-sample analysis on whole slide images (Fig. 3A-D,H-K). The immunofluorescence approaches (Fig. 2) required an optimized staining protocol with less unspecific background and tissue autofluorescence signal and several steps of semi-automatic and manual image analysis (see Materials and Methods section for details). In contrast, the automated deep learning-based approach (Fig. 3) was not limited to a region of interest, but was also used to analyse whole slide images containing multiple H&E-stained sections. Furthermore, this software tool directly obtained pixel-accurate values of individual fiber areas and surrounding endomysium thickness, eliminating the need for approximation, calculation, and correction of these primary parameters (see Materials and Methods section for details).

**Figure 2.**
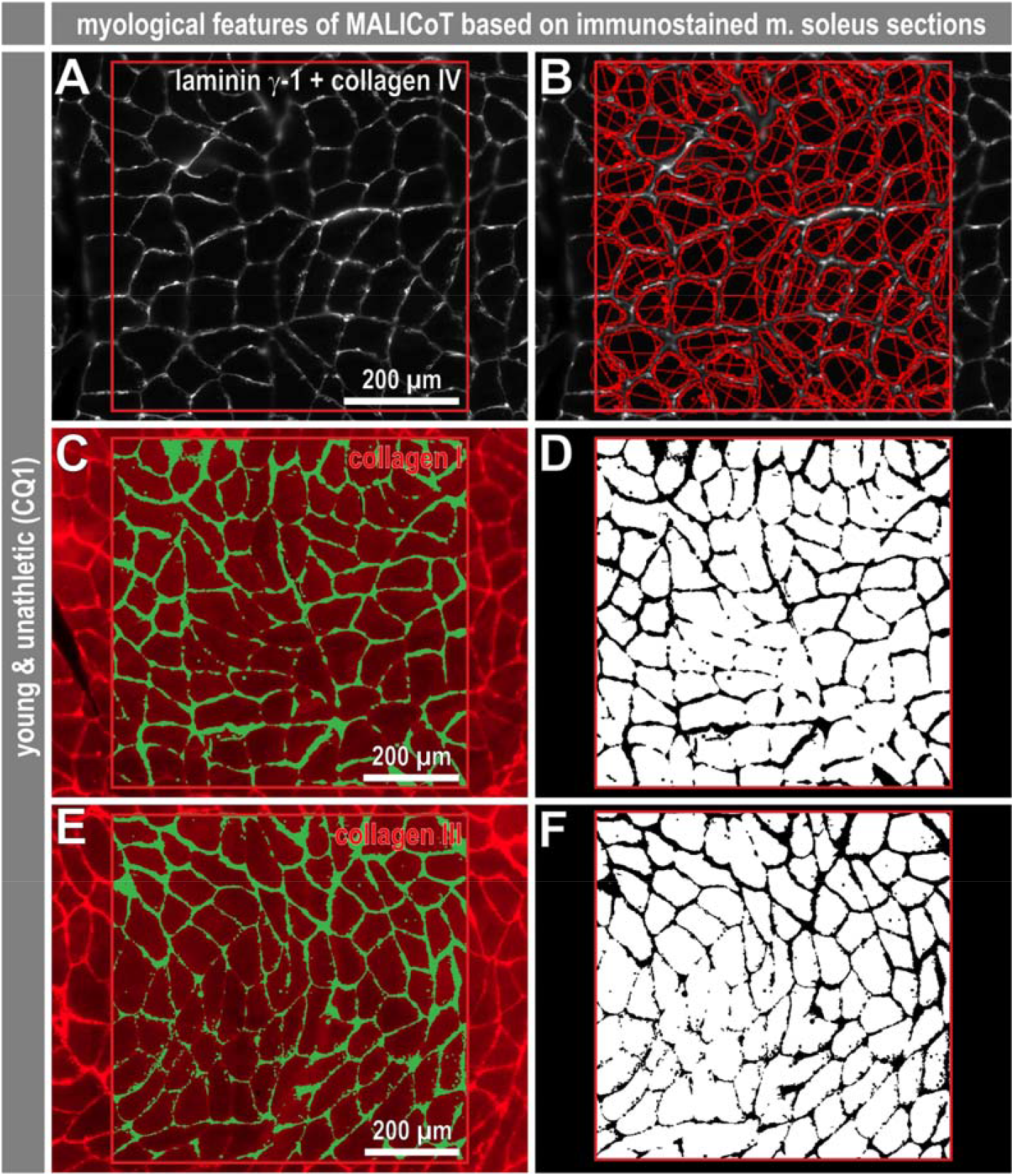
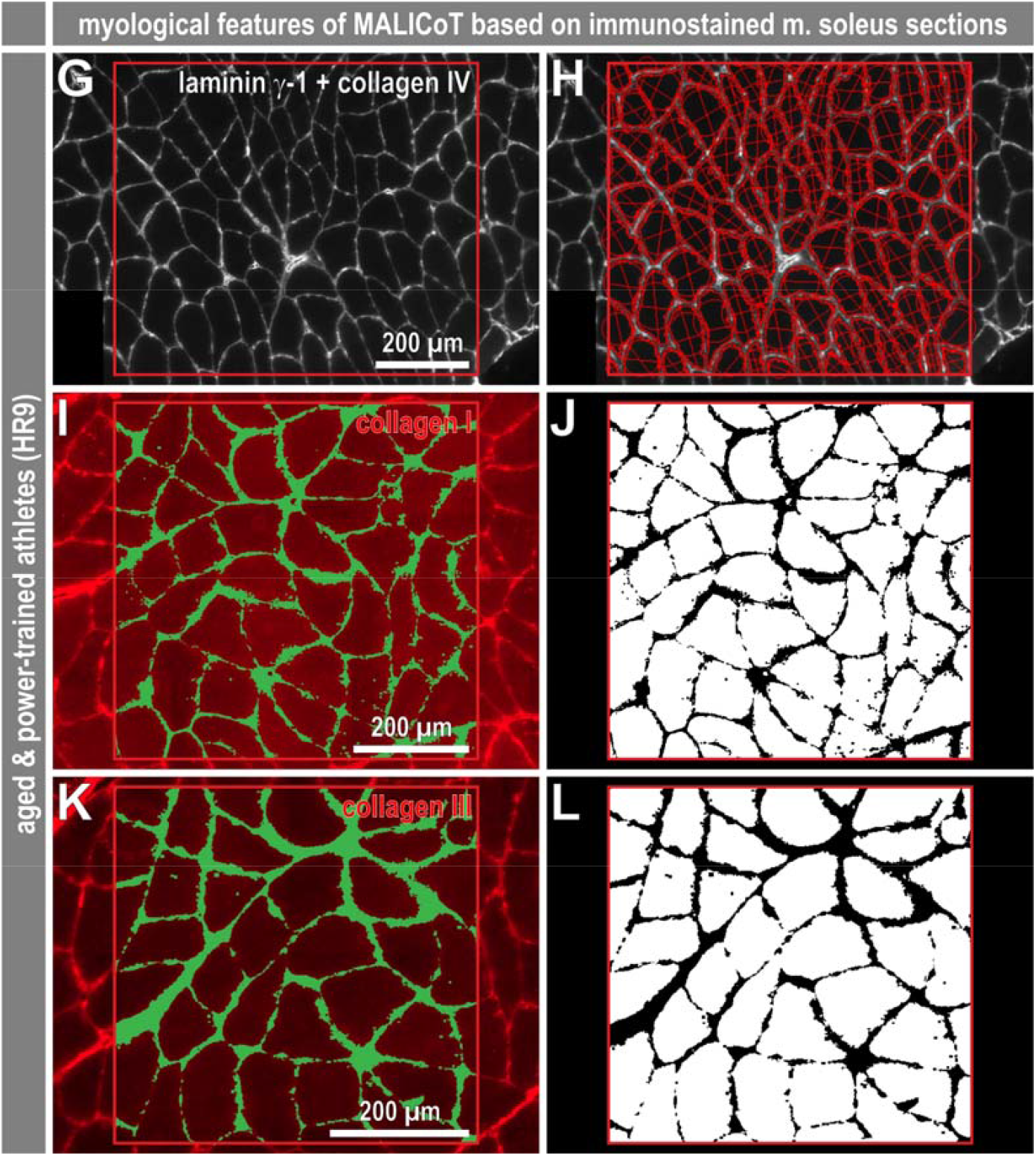
Myological parameters of MALICoT (I): immunostained sections of soleus muscle and manual digital image analysis. Exemplary images of soleus muscle sections from a young unathletic participant (**A, C, E**) and an aged athletic participant (**G, I, K**) double-immunostained for laminin-*γ*-1/collagen IV (**A, G**), and immunostained for either collagen I (**C, I**) or collagen III (**E, K**). The red squares indicate ROIs selected for segmentation of muscle fibers and endomysium using the microscope software ZEN. In the case of laminin-*γ*-1/collagen IV staining this analysis resulted in an approximation of individual fiber areas and Feret diameters (**B, H**) and a calculated determination of the endomysium. For collagen I and collagen III staining, the number of muscle fibers was counted and image analysis resulted in black and white images to determine the fiber (sum of white pixels) and endomysium (sum of black pixels) areas (**D, F, J, L**). Further details on analysis and calculation of derived parameters are provided in the Materials and Methods section.

**Figure 3.**
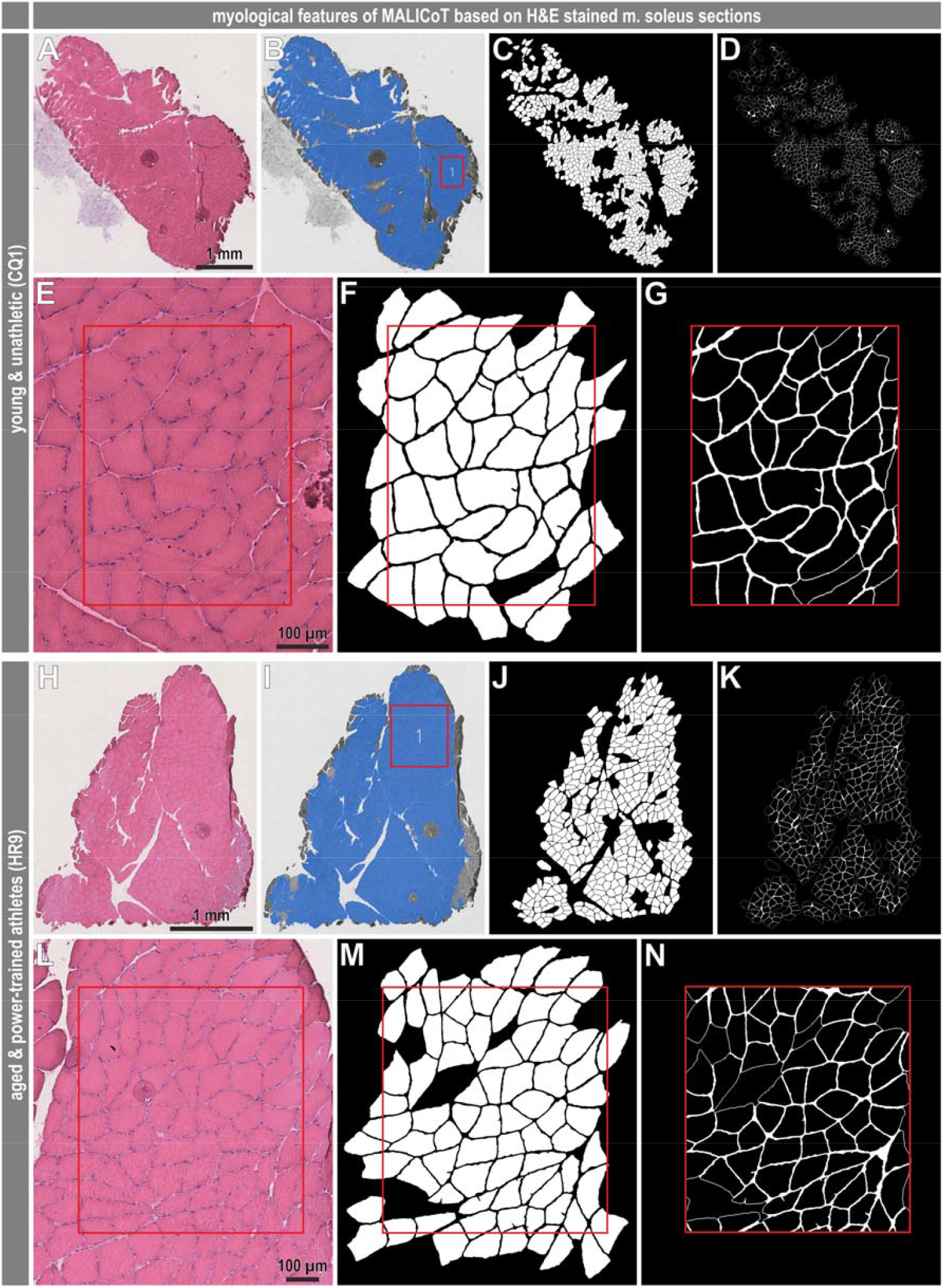
Myological parameters of MALICoT (II): H&E-stained sections of soleus muscle and automated deep learning-based image analysis. Exemplary images of H&E-stained soleus muscle sections from the same young unathletic (**A, E**) and aged athletic (**H, L**) participants as in figure 2. Muscle fibers (**C, F, J, M**) and endomysium (**D, G, K, N**) were automatically segmented for the whole sections (**A-D, H-K**) or ROI-based (red squares) (**E-G, L-N**). For whole section analysis, the sections and artefacts were automatically detected prior to segmentation (**B, I**), and areas of the sections that passed quality control parameters are shown in blue; for the ROI-based analysis, the ROI was manually set as in figure 2. Further details on image analysis and the quantitative parameters obtained are provided in the Materials and Methods section.

A comparison of the data obtained using the different quantification approaches showed similar overall results for the four main myological parameters shown, i.e., mean fiber area in µm^2^, mean endomysium thickness in µm, total endomysium area/total fiber area, and total endomysium area/fiber number in µm^2^ (Fig. 4). Automated deep learning-based analysis of H&E-stained soleus muscle sections in whole slide scans showed statistically significant increases in i) the mean fiber area with power-training in young participants and with aging in unathletic participants and ii) the total endomysium area/fiber number ratio also with power-training in young participants and with aging in unathletic participants, *and further between young unathletic participants and aged power-trained participants* (Fig. 4, row 1) (the italicised text is referred to in the following paragraph). The same approach showed very similar patterns in the scatterplots derived from the ROI-based analysis, where significant increases were detected for i) the mean fiber area with power-training in young participants *and with aging in unathletic* participants, ii) *the mean endomysium thickness between young unathletic participants and aged power-trained participants*, and iii) the total endomysium area/fiber number ratio with aging in unathletic participants (Fig. 4, row 2). ROI-based analyses of the laminin-*γ*-1/collagen IV, collagen I, and collagen III-stained soleus muscle sections showed statistically significant increases i) for laminin-*γ*-1/collagen IV and collagen III in the mean fiber area with power-training in young participants and with aging in unathletic participants, ii) for laminin-*γ*-1/collagen IV in the mean endomysium thickness and the total endomysium area/fiber number ratio with aging in unathletic participants, and iii) *for collagen I in the mean endomysium thickness with aging between athletic participants* (Fig. 4, rows 3-5). Two other significant changes should be considered with caution as they differ from the overall pattern in the other analysis approaches. The decrease shown for laminin-*γ*-1/collagen IV in the mean endomysium thickness with power-training in aged participants is contradicted by the *collagen I data showing an increase for this comparison* (Fig. 4, rows 3 and 4).

**Figure 4.**
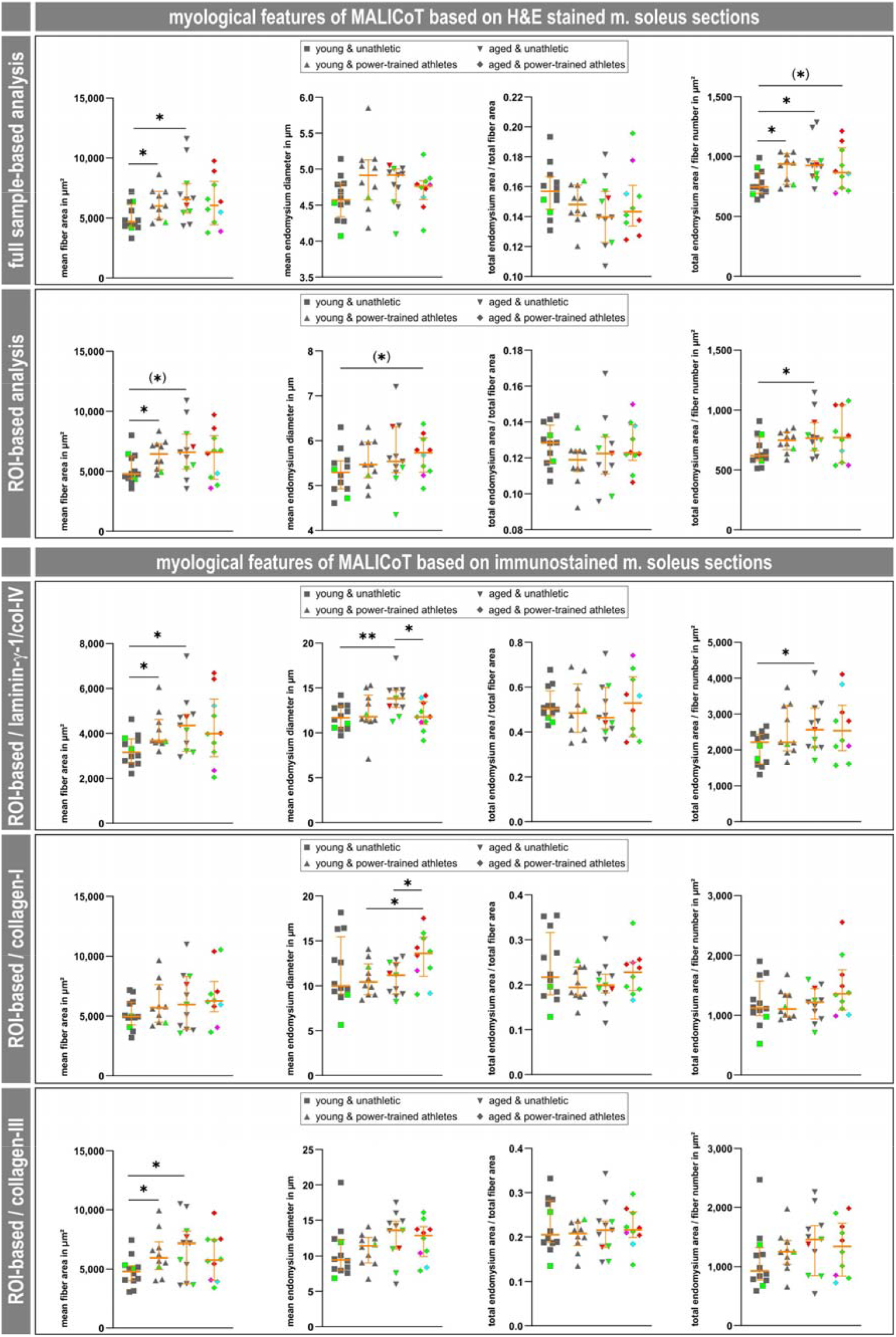
Compilation of myological parameters of MALICoT obtained through automated deep learning-based and manual digital image analysis of soleus muscle sections. Four main myological parameters are shown, i.e., mean fiber area in µm^2^, mean endomysium thickness in µm, total endomysium area/total fiber area, and total endomysium area/fiber number in µm^2^, for each of the five different digital image analysis procedures, i.e., H&E-stained full sample and ROI-based automated deep learning (rows 1 and 2) and fluorescence-stained extracellular matrix component ROI-based manual digital analysis (rows 3 to 5). In the case of automated analysis, the values for fiber area, endomysium thickness, total endomysium area and fiber number were obtained directly from the analysed images; in the case of the manual analysis, the values for the four parameters shown were derived from other primary parameters, depending on the type of manual analysis, from calculations using the values for total fiber area, total number of fibers, number of intact fibers, number of trimmed fibers, ROI dimensions, total fiber perimeter, or total endomysium area; details are provided in the Materials and Methods section. Each graph shows a scatter plot (young unathletic n=12, young power-trained athletic n=10, aged unathletic n=11, aged power-trained athletic n=10) with median and interquartile range (in orange). After normality testing, individual statistical significances were calculated using unpaired two-sample Welch’s t-tests, except for comparisons involving mean fiber area of young power-trained athletes from laminin/collagen IV staining, which were done using the Mann-Whitney U test, as this dataset did not have a normal distribution; p-values, **≤0.01, *≤0.05, (*)≤0.07. Dark grey symbols, normal skeletal muscle; green symbols, unspecific myopathological changes; red symbols, chronic neurogenic atrophy; magenta symbols, type II fiber atrophy; turquoise symbols, unspecific myositic changes; see Results section for details.

Next, the full data obtained using the different quantification approaches was adjusted for the biopsies with distinct myopathological diagnoses, i.e., only data from biopsies deemed to be normal or exhibiting unspecific myopathic changes was considered (Fig. 5). This reduced the number of biopsies from the group of aged unathletic participants by one into n = 10 and halved the number of biopsies from the group of aged athletes into n = 5. While the overall pattern of the scatter plots still remained very similar (Fig. 5), six group-specific comparisons lost statistical significance (indicated by the *italic text in the previous paragraph*). Analyzing the H&E-stained soleus muscle sections in whole slide scans still showed statistically significant increases in the mean fiber area with power-training in young participants and with aging in unathletic participants (Fig. 5, row 1), and in the total endomysium area/fiber number ratio also with power-training in young participants and with aging in unathletic participants (Fig. 5, row 1). The ROI-based analysis of H&E-stained sections solely showed a significant increase in the mean fiber area with power-training in young participants (Fig. 5, row 2). While there was no change in the ROI-based analysis of the laminin-*γ*-1/collagen IV- and collagen III-stained soleus muscle sections, the collagen I based analysis lost statistical significance (Fig. 5, rows 3-5).

**Figure 5.**
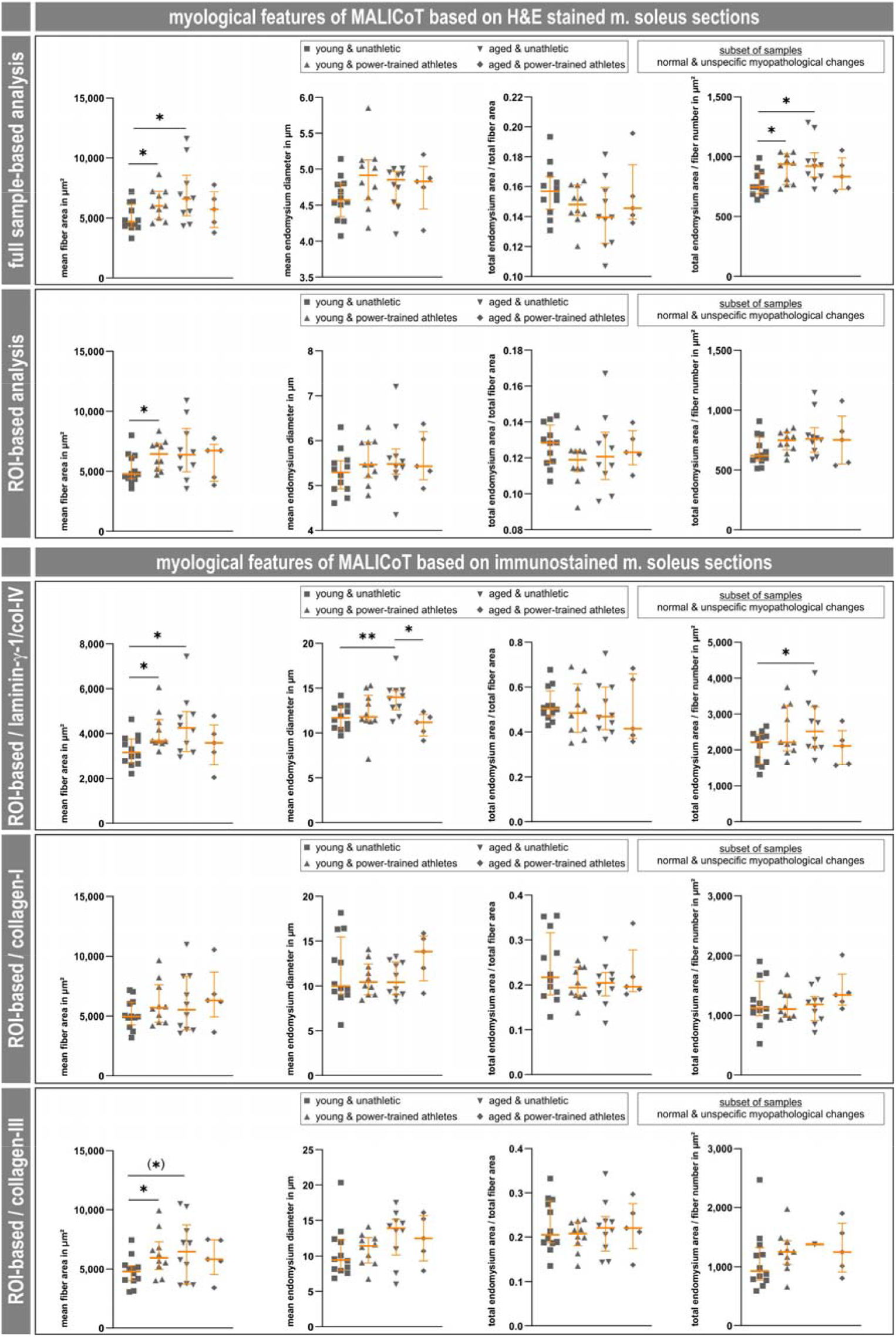
Adjusted MALICoT dataset excluding biopsies with distinct myopathological diagnoses. Four main myological parameters are shown, i.e., mean fiber area in µm^2^, mean endomysium thickness in µm, total endomysium area/total fiber area, and total endomysium area/fiber number in µm^2^, for each of the five different digital image analysis procedures, i.e., H&E-stained full sample and ROI-based automated deep learning (rows 1 and 2) and fluorescence-stained extracellular matrix component ROI-based manual digital analysis (rows 3 to 5). In contrast to figure 4, the compilation of myological parameters in this figure is based on the adjusted MALICoT dataset excluding biopsies with distinct myopathological diagnoses. Each graph shows a scatter plot (young unathletic n=12, young power-trained athletic n=10, aged unathletic n=10, aged power-trained athletic n=5) with median and interquartile range (in orange); the dark grey symbols in the scatter plots include both the biopsies with normal skeletal muscle and with unspecific myopathological changes. See the legend of figure 4 for additional details.

In summary, the following results were consistent and statistically significant: 1) A marked increase in the mean fiber cross-sectional area with power-training in young participants (5,026 ± 1,128 µm vs. 6,189 ± 1,298 µm, mean ± standard deviation, values related to the graphs in the first row of Fig. 5). 2) An even higher increase in the mean fiber cross-sectional area with aging in unathletic participants (5,026 ± 1,128 µm vs. 7,011 ± 2,441 µm). 3) Analogous increases in the ratio of total endomysium area to fiber number with power-training in young participants (777 ± 105 µm vs. 901 ± 122 µm) and with aging in unathletic participants (777 ± 105 µm vs. 952 ± 180 µm). Notably, no consistent and significant changes were found for the endomysium thickness or the ratio of total endomysium area to total fiber area.

### Proteomic analysis revealed decrease in plasma membrane calcium-transporting ATPase 2 in soleus muscle from aged power-trained athletes

In a next step, soleus muscle tissue samples from the 43 MALICoT participants were used for a proteotypic peptide-based proteomic analysis, resulting in the quantitation of 3,954 proteins (Tab. S1). The principal component analysis (PCA) revealed a picture of individual proteomes, but no grouping in terms of the MALICoT participant classification (Fig. 6A). As the heat-map plot of the proteomic data from the four MALICoT groups also showed no sample clustering, further standard data analysis was omitted. In addition, manual inspection of the data showed that there were no changes in the abundance of any of the detected collagens, laminins, nidogens, or other proteins related to the composition of intramuscular connective tissue such as dermatopontin, fibrillin-1, and perlecan (basement membrane-specific heparan sulfate proteoglycan core protein) (Tab. S1). However, in a proteomic dataset, changes for which no fold change can be calculated may also be of interest. Inspection of the dataset for proteins that were detected in some samples but below the detection limit (or not expressed at all) in other samples of the MALICoT groups led to the identification of a single candidate of interest, plasma membrane calcium-transporting ATPase 2 (ATP2B2 or PMCA2). While plasma membrane calcium-transporting ATPases 1 and 4 (ATP2B1 and ATP2B4) were detected in all samples from all four MALICoT groups, ATP2B2 was detected in 12 out of 12 samples from young unathletic participants, in 9 out of 10 from young power-trained athletes (not detected in sample QC6) and in 9 out of 11 from aged unathletic participants (not detected in samples MS7 and UI2), but only in 2 (samples DP4 and XV3) out of 10 from aged power-trained athletes (Fig. 6B). To verify this marked difference in the abundance of ATP2B2, the proteomic analysis was complemented by ATP2B2 immunofluorescence imaging of samples from aged unathletic participants and aged power-trained athletes. In contrast to a spot chain-like sarcolemmal localisation of the ATP2B2 signals in the samples from aged unathletic participants, no such signal enrichment was found at the membrane in samples from aged power-trained athletes (Fig. 7).

**Figure 6.**
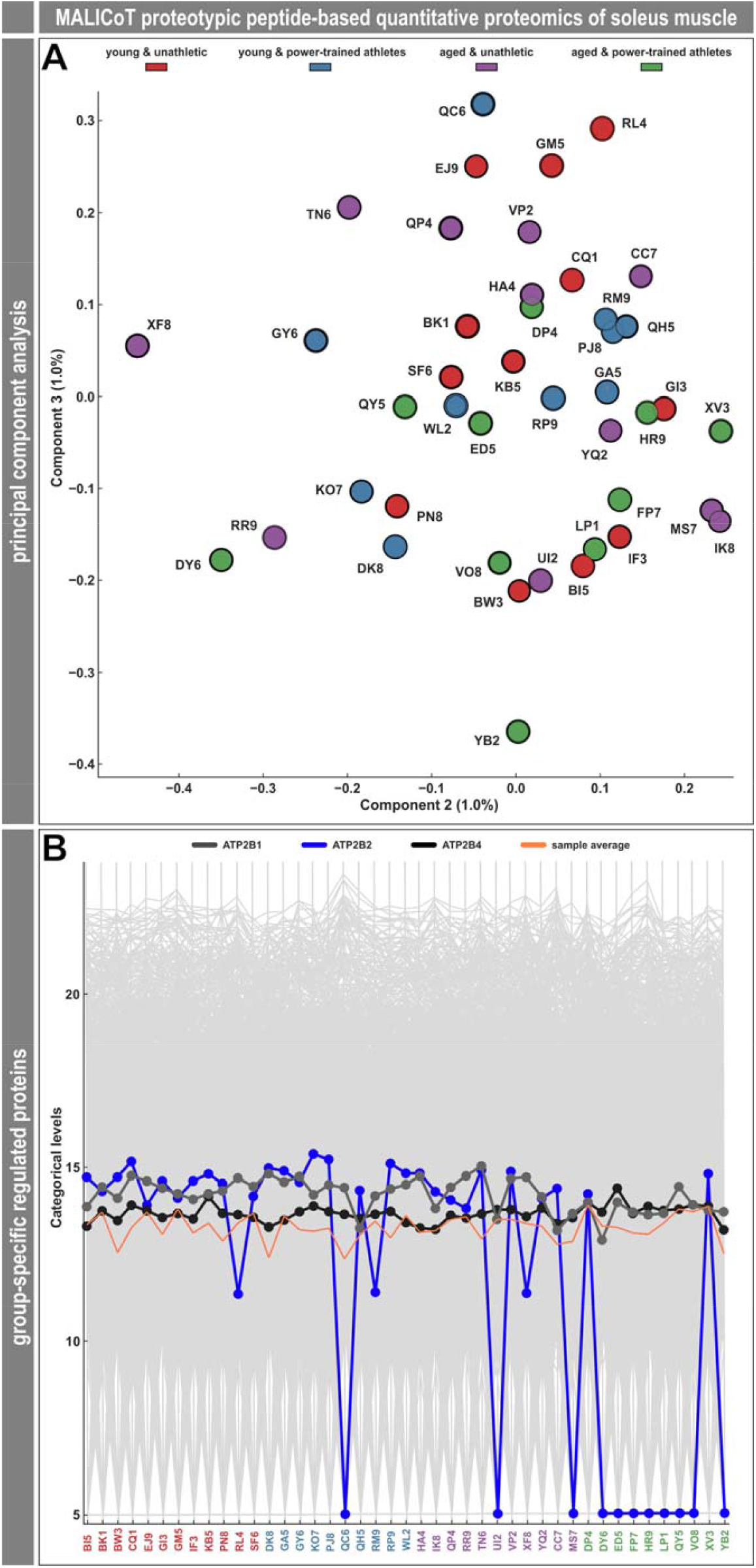
Proteotypic peptide-based quantitative proteomic analysis of MALICoT-derived soleus muscle tissue. (**A**) Based on the quantitation of 3,954 proteins (Tab. S1), principal component analysis (PCA) determined that there were no major differences in the global protein expression pattern. As the samples from the four participant groups could not be grouped by principal component analysis and clustering, further global data analysis was omitted. (**B**) Manual inspection of several muscle and connective tissue-related proteins revealed similar abundances in all samples. However, one of the quantitated proteins, plasma membrane calcium-transporting ATPase 2 (ATP2B2), was not detected in 8 out of the 10 samples from the aged power-trained athletes. Color code as in figure 1 and supplementary table 1.

**Figure 7.**
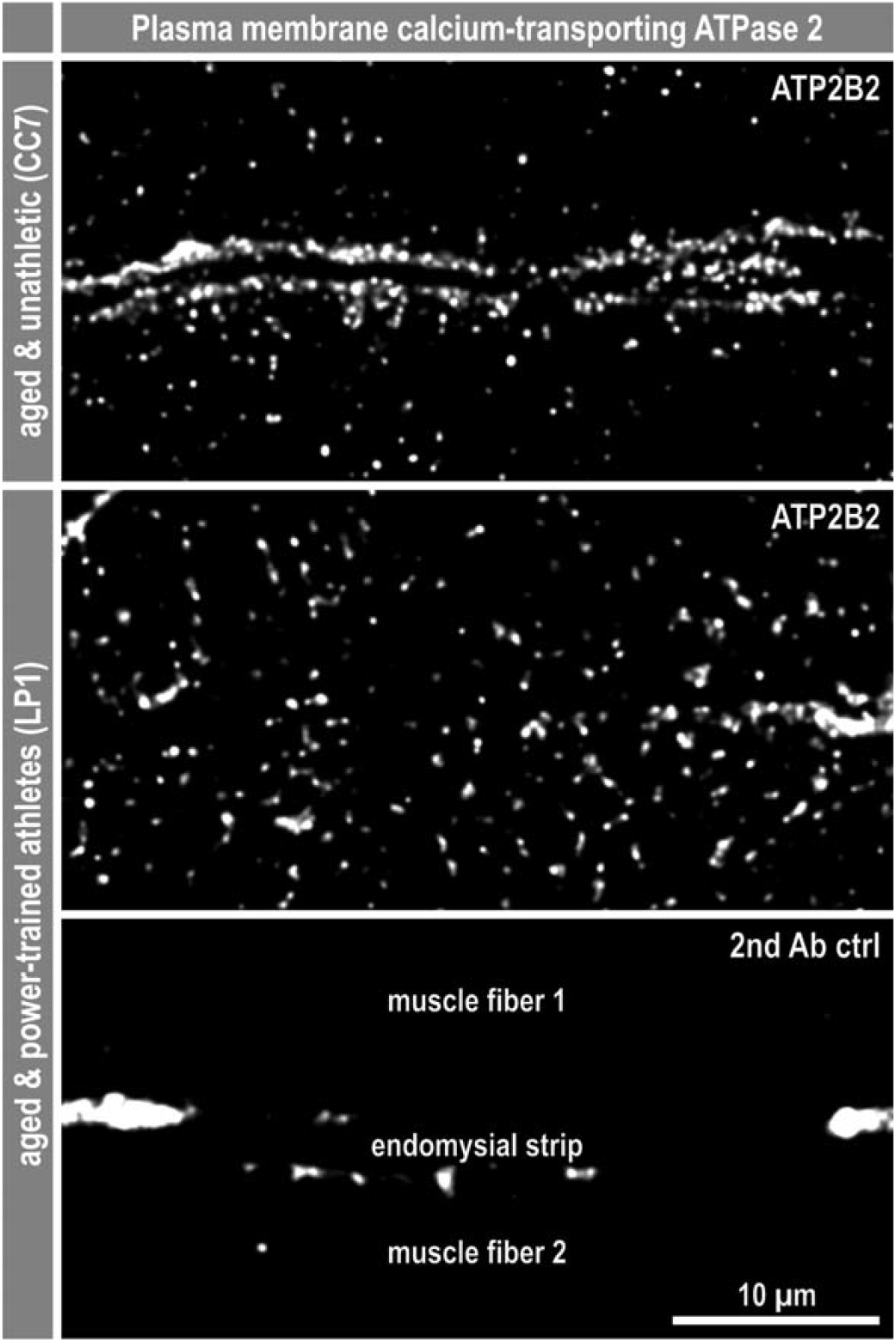
Aged power-trained athletes have less plasma membrane calcium-transporting ATPase 2 at the sarcolemma. To verify the proteomic finding of a reduction of plasma membrane calcium-transporting ATPase 2 (ATP2B2 or PMCA2) below the detection limit specifically in the aged power-trained athletes, samples from aged unathletic (HA4, TN6, YQ2, CC7) and aged power-trained athletes (HR9, LP1, VO8, YB2), who had no specific diagnosis (Fig. 1), were immunostained for ATP2B2. (**A**) As expected, the subcellular distribution of ATP2B2 in the aged unathletic samples appeared as strings of spots in a row at the level of the two sarcolemmas in the center of the field of view. (**B**) In contrast, there was no signal enrichment at the sarcolemma in samples from aged power-trained athletes, but only single spots in the sarcoplasm. (**C**) Negative controls with secondary antibody only show few artefacts from tissue autofluorescence and no ATP2B2-specific signals. All three images show a section of sarcoplasm from one muscle fiber, the endomysium in the center of the image, and a section of an adjacent muscle fiber below. The confocal images were deconvolved using Huygens Essential (Scientific Volume Imaging B.V., Hilversum, The Netherlands).

## Discussion

The Master Athletic Laboratory Study of Intramuscular Connective Tissue (MALICoT, DRKS00015764) primarily set out to determine the endomysium content in the human soleus muscle as a function of athletic exercise and age. Based on previously published results, it was assumed that aging (Alnaqeeb et al., 1984; Fede et al., 2022; Yeung et al., 2023) and low physical activity (Thot et al., 2021) might lead to increased endomysium content. To the best of our knowledge, no studies addressing endomysium content as a function of age and training state have been published in humans. Comparing Masters athletes with sedentary individuals at different ages thus allows to study age-related muscle changes without the confounding factor of sedentary behavior (Rittweger et al., 2004). Another key aspect of this work was to compare our previously published approach of quantifying endomysium content and muscle fiber size using laminin-*γ*-1 immunostaining of muscle tissue cryosections ((Thot et al., 2021), in this work conducted as laminin-*γ*-1/collagen IV double-immunostaining) with collagen I and collagen III immunostainings, as well as deep learning-based artificial intelligence analysis of H&E-stained cryosections (Mill et al., 2025).

### Deep learning-based analysis of H&E-stained muscle sections is a new state of the art technique

While laminin-*γ*-1 and collagen IV antibodies stain the basement membrane and delineate the endomysial space, allowing segmentation of muscle fibers in immunofluorescence images (Fig. 2A,B,G,H), the endomysium content must be calculated (Thot et al., 2021). Conversely, when using antibodies directed against collagen I and collagen III, the endomysium area is labelled (Fig. 2C-F,I-L) and the muscle fiber cross-sectional area must be derived. All immunostaining variants require calculation, and to a certain extent the estimation, of fiber area and endomysium thickness. To do so, manual work is required with regard to immunofluorescence staining protocols, immunofluorescence microscopy and image processing. In contrast, the deep learning-based image analysis only requires standard H&E-stained muscle tissue cross-sections and can also work with whole slide images containing multiple stained sections acquired by a slide scanner (Fig. 3). Most importantly, this approach obtains pixel-accurate values of fiber area and endomysium thickness from ROIs or single and multiple sections (Mill et al., 2025).

Overall, the scatter plot patterns (Fig. 4) appeared similar and consistently showed statistical significance between groups for the mean fiber cross-sectional area and the total endomysium area to fiber number ratio obtained by the deep learning-based analysis (Fig. 4, rows 1 and 2), as well as for the laminin-*γ*-1/collagen IV and collagen III immunostains (Fig. 4, rows 3 and 5). Bland-Altman plots of the two parameters, mean fiber area and total endomysium area to fiber number ratio, revealed agreement between ROI-based analysis of H&E-stained and laminin-*γ*-1/collagen IV immunostained sections. In the case of mean fiber area, there was positive simple proportional bias (Fig. 8A), and in the case of total endomysium area to fiber number ratio there was negative simple proportional bias (Fig. 8B). Additional Bland-Altman plots showed consistency between the whole slide scan-based and the ROI-based analyses of H&E-stained soleus muscle sections. In the case of mean fiber area, there was close to zero mean difference (Fig. 8C), and in the case of total endomysium area to fiber number ratio, there was positive mean difference (Fig. 8D). The latter can be explained by the fact that the ROIs were set to regions of the muscle cross-sections that did not contain perimysium. Since the segmentation masks derived from the deep learning-based analysis precisely fit the H&E staining pattern ((Mill et al., 2025); and this work), and the Bland-Altman plots demonstrated that the deep learning-based analysis of H&E-stained muscle sections is an appropriate and simple substitute for the immunostaining approaches, this approach should be considered state of the art technique.

**Figure 8.**
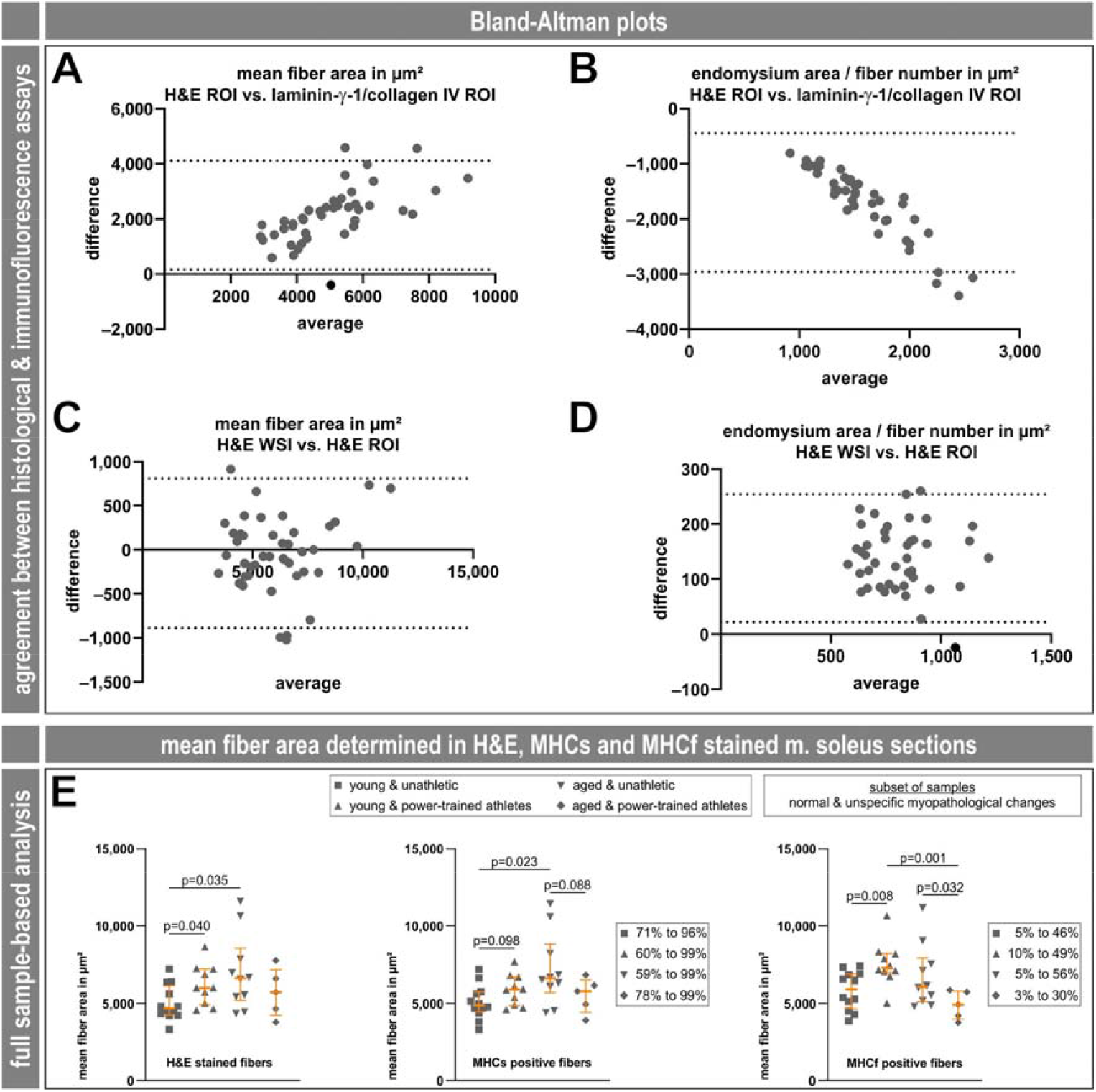
Bland-Altman plots of agreement between analysis approaches, and attribution of the significantly increased mean fiber area to the fiber type. (**A**-**D**) Bland-Altman plots of mean fiber area and total endomysium area to fiber number ratio for ROI-based analysis of H&E-stained and laminin-*γ*-1/collagen IV immunostained sections as well as of whole slide scan-based and ROI-based analyses of H&E-stained soleus muscle sections. (**E**) Scatter plots with median and interquartile range (in orange) of the mean fiber area in µm^2^ derived from full sample automated deep learning analysis of H&E (same graph as shown in figure 5, but with exact p-values), MHCs and MHCf stains, limited to the diagnoses-adjusted MALICoT dataset. For MHCs and MHCf, the range of positively stained fiber fractions as a percentage is indicated.

### Power-training and aging have no significant impact on the endomysium content of human soleus muscle

In rat soleus and extensor digitorum longus muscles, a marked increase in endomysium and collagen content with age has been reported, which correlated with increased muscle stiffness (Alnaqeeb et al., 1984). In mouse lateral gastrocnemius muscle, it was found that the abundance of multiple extracellular matrix proteins in the intramuscular connective tissue increased with age, while wheel running exercise had no effect (Yeung et al., 2023). In human vastus lateralis muscle, endomysium content and collagen staining intensities were basically similar between endurance-trained athletes and untrained controls (Mackey et al., 2005). In general, it seems that there is little published data on the relationship between intramuscular connective tissue or endomysium content and physical activity or age, and the results are contradictory.

Based on the complete MALICoT data set, neither the mean endomysium thickness nor the ratio of endomysium to muscle fiber area changed significantly with athletic exercise or age (Fig. 4, the two middle graphs per row). Only the ratio of the endomysium area to fiber number increased with power-training in young individuals and with aging (Fig. 4, the right graph per row). Simultaneously, a destructive interaction (which causes less effect) seemed to appear between athletic exercise and age (Fig. 4, top right graph). However, looking at the results from the adjusted MALICoT dataset that excluded biopsies with distinct myopathological diagnoses (Fig. 5), the latter interaction was no longer present.

The additional proteomic analysis also revealed no significant changes in the abundance of proteins related to the composition of intramuscular connective tissue (Tab. S1). The only proteomic finding, which was also confirmed by immunofluorescence analysis, was a marked reduction of plasma membrane calcium-transporting ATPase 2 (ATP2B2 or PMCA2) in this group of aged power-trained athletes (Figs. 6B, 7). No data is currently available on the function of this ATP-dependent Ca^2+^ pump isoform in skeletal muscle cells (Stauffer et al., 1993, 1994). ATP2B2 (UniProt entry Q01814) may be involved in regulating the basal sarcoplasmic Ca^2+^ concentration or transporting Ca^2+^ to the endomysial space.

### Power-training and aging significantly increase the fiber cross-sectional area in human soleus muscle

The complete and the diagnoses-adjusted MALICoT datasets showed a significant increase in the mean fiber cross-sectional area with power-training in young individuals and even more prominent with aging (Figs. 4 and 5, top left graph in each). The finding of an increased mean fiber area with power-training in human muscle is well-documented and confirms previously reported findings (for example, (Häggmark et al., 1978; Hermansen and Wachtlova, 1971). However, the influence of aging on fiber cross-sectional area in human muscle is apparently complex and inconsistent. Here, a meta-analysis including 19 different studies on male vastus lateralis muscle and 1 on male gastrocnemius muscle (figure 3 and table 3 in reference (Lee et al., 2024) revealed that the fiber cross-sectional area can markedly decrease, remain approximately the same, or increase markedly with age. The few studies on female muscles included in this meta-analysis showed the same pattern (Lee et al., 2024). To include the fiber type in our finding of a significant increase in the fiber cross-sectional area with power-training and aging, we additionally analysed the MHCs- and MHCf-stained sections using the deep learning-based image analysis software. In the diagnoses-adjusted MALICoT data, the soleus muscle contained a high proportion of MHCs-positive type I fibers (between 59% and 99%, as expected) and lower amounts of MHCf-positive type II fibers (between 3% and 56%) (Fig. 8E). For both fiber types the patterns of mean fiber area changes between the four groups were basically the same as that detected by H&E staining, however, with different levels of statistical significance (Fig. 8E). In this respect, MALICoT adds first data on human male soleus muscle.

Notably, when considering such significant changes in the fiber area (or ‘fiber cross-sectional area’, not to be confused with ‘muscle cross-sectional area’) and the fact that muscle tissue specimens of essentially constant dimensions are analyzed by means of a muscle biopsy, a change in values referring to the fiber number is inevitable. In case of MALICoT, the significantly increased fiber area associated with a decreased fiber number in the cross-sections of the biopsy specimens can explain the increase in the endomysium area-to-fiber-number ratio. Therefore, the change in this relative value has to be considered as an artifact and should not be considered further.

In summary, MALICoT, with evidence from a group size of roughly 11 male participants, showed as primary result that athletic exercise and age did not significantly affect endomysium thickness or area in the human soleus muscle. Since 17 of the 43 healthy participants exhibited relevant histopathological findings, a detailed myopathological analysis seems necessary for studies examining the effects of age or exercise, even when participants are asymptomatic. Additionally, this study revealed that the fiber cross-sectional area in the human soleus muscle increased significantly with athletic exercise and age.

## Data availability

Proteomic raw data have been deposited to the ProteomeXchange Consortium (Deutsch et al., 2023) via the PRIDE (Perez-Riverol et al., 2022) partner repository with the dataset identifier PXD070244. Anonymized subject data used and analysed for this report will be shared upon reasonable request.

## Acknowledgments

We gratefully acknowledge the dedicated contribution of the study participants.

## Grants

This research received no specific grant from any funding agency in the public, commercial, or not-for-profit sectors.

## Disclosures

The authors declare that they have no competing interests. C.S.C. and R.S. serve as consultants for MIRA Vision Microscopy GmbH.

## Ethical approval

This study was approved by the Ethics Committee of the North Rhine Medical Association in Düsseldorf (#2018269), Germany, and registered in the German Clinical Trials Register (DRKS-ID DRKS00015764). All participants gave written informed consent before participating in the study.

## Author contributions

C.S.C., R.S., and J.R. conceived and designed research. C.S.C., S.W.H., C.B., A.S., and J.R. performed experiments. All authors analyzed data. All authors interpreted results of experiments. C.S.C. prepared figures. C.S.C. and J.R. drafted the manuscript. C.S.C., R.S., and J.R. edited and revised the manuscript. All authors, except for J.R. who died before (Frings-Meuthen et al., 2025), approved the submitted version of the manuscript.

## Table legends

**Supplementary Table 1. Results of the proteome analysis**. Data set from proteotypic peptide-based quantitative proteomic analysis using soleus muscle tissue samples from the 43 MALICoT participants. Mean log_2_ intensities of 3,954 proteins; color code as in figures 1 and 6.

## Notes

https://www.ebi.ac.uk/pride/archive/projects/PXD070244

### Summary of Updates

Proteomic dataset identifier added. Quantitative data of MHCs and MHCf stains added (figure 8E); manuscript text updated accordingly.

